# Discovery of Prognostic Biomarkers in Gastric Cancer Through Machine Learning and Bioinformatics Analysis of Gene Expression Data

**DOI:** 10.1101/2025.07.29.667579

**Authors:** Rohit Kumar Verma, Prashant Kumar Srivastava, Ashutosh Singh

## Abstract

Gastric cancer (GC) often gets diagnosed in its advanced stages, resulting in poorer prognoses. To identify potential biomarkers in GC, we used fifteen datasets from NCBI-GEO and integrated them into a complete dataset. From the complete dataset, we extracted a subset of only the known cancer driver genes. Using Recursive Feature Elimination (RFE), mutual information (MI), and tree-based (TB) method (SelectFromModel), we extracted top gene features using RFE (10, 20, 30, 40, and 50) and 50 features each from MI and TB method. Subsequently, we applied machine learning classifiers to these selected gene features to classify cancer and normal samples. The SVC classifiers demonstrated better performance when utilizing the top 50 gene features using RFE and MI, while the AB classifier achieved the highest performance using TB for the complete dataset and for driver datasets, RF performed well using 40 features using RFE and SVC, and ET showed up better performance using 50 features using MI and TB feature selection for the test dataset. After combining all the genes from both datasets from and from all three feature selection methods, only 115 showed differentially expressed genes. A Lasso-penalized Cox regression model was applied to narrow down the gene selection to fourteen. This study highlights the effectiveness of integrating machine learning and bioinformatics analysis to identify new biomarkers for GC.

## 1. Introduction

Gastric cancer (GC) ranked as the fifth most prevalent cancer and stood as the fifth principal contributor to cancer-related fatalities globally in the year 2022 [1]. Approximately 1.1 million new cases of GC were projected, leading to an estimated 770,000 deaths in the same year. The incidence rates displayed an average twofold increase in males compared to females, with rates of 15.8 per 100,000 and 7.0 per 100,000, respectively. Variations were observed across countries [2]. Despite notable advancements in the survival rates of individuals with GC over recent decades, the diagnosis of GC frequently occurs at an advanced stage, and prognoses remain unsatisfactory. This is primarily attributed to the elevated recurrence rates of the condition [3]. Optimal approaches for lowering mortality involve interventions focused on early detection, systematic prevention, and personalized therapy. Meanwhile, conventional treatment methods like surgery may have reached a limit in terms of controlling the disease locally and reducing mortality. This underscores the challenge that GC continues to be inadequately curable on a global scale [4].

GC is characterized by its heterogenous, with each patient presenting a unique genetic and molecular makeup [5]. Despite extensive research on molecular biomarkers, many identified ones have not stood up to validation studies. Although various biomarkers are available for GC, carcinoembryonic antigen (CEA) and CA19-9 remain the primary biomarkers employed in clinical practice for diagnosing GC [6–7]. Other conventional biomarkers like CA72-4, alpha-fetoprotein, and CA125 have been extensively used in diagnosing GC [8–10].

Driver mutations are common in GC; however, mutations have been identified in a range of genes, such as TP53, SYNE1, CSMD3, LRP1B, CDH1, PIK3CA, ARID1A, PKHD, KRAS, JAK2, CD274, and PDCD1LG2 [11–13]. Elevated tumor mutational burden (TMB) could indicate a positive outlook following surgical removal of GC tumors, yet it may lead to reduced effectiveness of postoperative chemotherapy or chemoradiotherapy, specifically within stage Ib/II cases. This phenomenon might be influenced by TMB-related immune cell infiltration and hypoxia, respectively [14]. Presently, GC has few targetable or actionable mutations known so far, which makes it challenging to treat effectively. Therefore, the identification of new biomarkers with clinical significance is imperative. This will not only assist in diagnosis but also improve the prognosis for patients with GC.

The different ML approaches have been studied earlier, using miRNA data to identify biomarkers. Gilani et al., 2022 identified hsa-miR-1343-3p as a promising candidate for investigation as a diagnostic biomarker for GC, employing the Boruta feature selection technique and a machine learning approach [15]. Chen et al., 2022 identified 3 hub genes mainly ATP4A, ATP4B, and ESRRG which are strongly associated with GC. However, by utilizing the ML models for GC with different supervised learning models, they achieved a higher AUC value (0.986) [16]. A similar study has been done by Azari et al., 2023 discovered that hsa-miR-21, hsa-miR-133a, hsa-miR-146b, and hsa-miR-29c were linked to increased mortality and could potentially help in the early detection of GC patients. Through machine learning, they identified a set of dysregulated miRNAs that could serve as potential biomarkers for GC [17]. Thus, Next-generation sequencing (NGS) is vital in evaluating the molecular condition of individuals with cancer. It enables thorough examination of all human genes, aiding in identifying the actionable driver mutations suitable for targeted therapeutic approaches [18].

Our research involved a comprehensive integration of machine learning techniques and bioinformatics analysis to identify potential genes associated with GC, with the aim of establishing them as promising prognostic indicators. We analyzed fifteen datasets obtained from microarray studies to discern genes that effectively and accurately classifying between GC and normal tissue samples, utilizing ten different supervised machine learning models for classification. Furthermore, we also conducted a comparison analysis of the gene signatures obtained from ML with those of DEGs. Using the DEGs, we constructed a LASSO Cox regression model based on data from TCGA-STAD, which enabled the identification of genes with prognostic potential. Additionally, we also conducted survival analysis, WGCNA, and pathway analysis with the obtained prognostic biomarkers.

## 2. Materials and Methods

### 2.1. Microarray datasets download and pre-processing

The CEL files of the gene expression profiles of fifteen independent GEO datasets for GC, which comprised 1090 cancer and 310 normal samples, were downloaded from the NCBI GEO database (https://www.ncbi.nlm.nih.gov/geo/) and TCGA-STAD (https://portal.gdc.cancer.gov/) to conduct survival analysis on potential prognostic genes [19]. We obtained fifteen datasets from the same platform (GPL570) using the hgu133plus2 annotation package. Using frozen Robust Multiarray Averaging (fRMA), the dataset was background corrected, normalized, and log2 transformed for further downstream processing. Further, the datasets were mapped probe to gene mapping using microarray dependent platform to gene symbol. The Entrez IDs were excluded for genes lacking corresponding gene symbols [20]. The samples from NCBI-GEO utilized in our study are provided in **Supplementary Table S1.** Fifteen datasets were combined using the R cbind function, wherein they were merged based on gene symbol, followed by undergoing expression normalization to ensure consistent expression distribution [21].

### 2.2. Batch-effect removal and dataset conversion Center effect

Integrating multiple datasets into one can introduce batch effects or variations that are not biologically relevant. To overcome this issue, adjustments were executed utilizing the Empirical Bayes algorithm via the ComBat function within the sva library in R [22]. Subsequently, probes with low variant probes were filtered out from the Meta-dataset to aid in machine learning analysis and differential expression analysis. To validate the effectiveness of batch-effect correction on the dataset transformed using ComBat, Uniform Manifold Approximation (UMAP) was plotted in R, indicating the simplification of complex data. Following preprocessing and batch-effect correction, the merged dataset yielded an initial compilation consisting of 1400 samples and 20,161 gene features, which will be termed as the complete dataset [23]. Additionally, for datasets pertaining to cancer driver genes, we utilized three databases such as Bailey, COSMIC, and IntOGen, as previously discussed by Joon et al., 2023 for extracting the cancer driver genes [24–27]. We merged cancer-driver gene lists from three databases, extracting unique genes, yielding 1304 known cancer-driver genes.

### 2.3. Feature Selection and Training Classification Model

After removing the batch effect, the dataset underwent training of supervised machine learning algorithms, which were applied to both the entire datasets and subsets containing cancer driver genes. Feature selection techniques play a critical role in gene expression datasets, primarily because of their high dimensionality. In such datasets, the number of genes typically far surpasses the number of samples available. These methods serve to identify the most important genes that play a significant role in the biological processes or outcomes being studied while filtering out irrelevant or redundant ones. By trimming down the dimensionality and concentrating on essential features, feature selection aids in enhancing interpretability, cutting down computational complexity, and boosting the predictive accuracy of machine learning models applied to gene expression data. This leads to more precise and reliable analyses in both biological research and clinical settings. First, we used Pearson’s correlation coefficient algorithm to eliminate surplus features from the complete dataset. Next, we applied AUC-based feature selection on the complete dataset. Finally, we used two from the wrapper and one from the filter method of feature selection, i.e., Recursive feature elimination (RFE), tree-based (SelectedFromModel), and Mutual information (MI) algorithms to identify the most important features [28–31]. A detailed methodology for feature selection is outlined in the **Supplementary Methodology sections 1.1 and 1.2.**

### 2.4. Differential Gene Expression Analysis

We’ve also incorporated GTEx normal samples in the expression analysis for GC alongside both cancerous and normal samples [32–33]. A comprehensive methodology for expression and statistical analysis is provided in **Supplementary Methodology, Section 1.3.** To identify differentially expressed genes, we utilized the complete dataset for DE gene analysis. We employed the limma package and applied standard statistical criteria (logFC ≥ 1 and logFC ≤ −1 and p < 0.05) to identify DE genes in the dataset [34]. Subsequently, the machine learning (ML) obtained gene features were compared with the DE genes, and the overlapping genes underwent further analysis to elucidate the pathways involved, utilizing a chord diagram using SangerBox, which is an online user-friendly database [35].

### 2.5. Survival analysis

The intersection of genes found among DEGs and the features derived from both the complete and driver genes were subjected to refinement through Lasso-penalized Cox regression. The survival analysis was conducted using the TCGA-STAD expression dataset by categorizing patients into high and low-risk groups based on the expression levels and multivariate Cox regression coefficients. Survival analysis was performed using KM plotter, an open-source web-based tool (https://kmplot.com/analysis/) [36–37]. Samples from patients were sorted into high and low mRNA expression categories for individual genes to examine overall survival. A lasso-penalized Cox regression analysis was also employed to confirm the relationship between the predictive genes and patient survival outcomes. A significance threshold of p < 0.05 was utilized for both survival and Cox regression analyses. [38].

### 2.6. Construction of co-expression network (WGCNA) and identification of key module genes

To determine if the genes identified through Lasso penalized Cox regression are not only differentially expressed but also coexpressed with other genes, we used the WGCNA package to construct a scale-free coexpression network based on the TCGA-STAD dataset [39]. We determined the soft-thresholding parameter β for a free-scale network. To thoroughly analyze functional modules, the adjacency matrix was transformed into a topological overlap matrix (TOM). Subsequently, we calculated the dissimilarity matrix between genes (dissTOM = 1 - TOM). Hierarchical clustering of dissTOM grouped genes with similar expression into distinct gene modules. The DynamicTreeCut algorithm was then employed to identify these gene modules, and modules with high similarity were further merged. Module eigengenes (MEs) were utilized to identify gene modules associated with tumor and normal tissues. MEs, representing the core element of each gene module, were used to summarize the expression patterns of all genes within a specific module. We then calculated the correlation between tumor, normal samples, and MEs to identify clinically significant modules. Using the Fisher exact test, we identified the significant modules, for the fourteen genes which we identified through Lasso-Cox penalized regression. The blue, red, and turquoise modules emerged as significant modules; however, the red module did not demonstrate clinical relevance. To further investigate the pathways associated with this module, we performed KEGG pathway enrichment analysis using ShinyGO v0.77 on the significant module identified [40–42].

## 3. Results

### 3.1. Gene panels distinguishing gastric cancer and normal tissue samples

Our main goal is to identify potential markers by integrating various datasets from microarray studies in GC. The complete dataset was merged using ComBat to remove batch effects. The boxplot indicates distinct differences in the sample distributions of each dataset before batch effect removal, implying the presence of such an effect. Following batch effect removal, the distributions appear more consistent, aligning their medians along a common line **Figure 1 (A-B)**. Similarly, we have shown the distribution of samples before and after applying the ComBat as density plot **Figure 1 (C-D)**. The UMAP diagram reveals the initial clustering of samples from each dataset, indicative of the batch effect. However, post-removal, samples from different datasets exhibit interwoven clusters, suggesting successful removal of the batch effect **Figure 1 (E-F)**. Collectively, these findings support the efficacy of batch effect removal. This consolidation resulted in 20,161 genes consistently present across all datasets with identical gene symbols. Additionally, cancer driver genes comprise 1174 genes.

**Figure 1:**
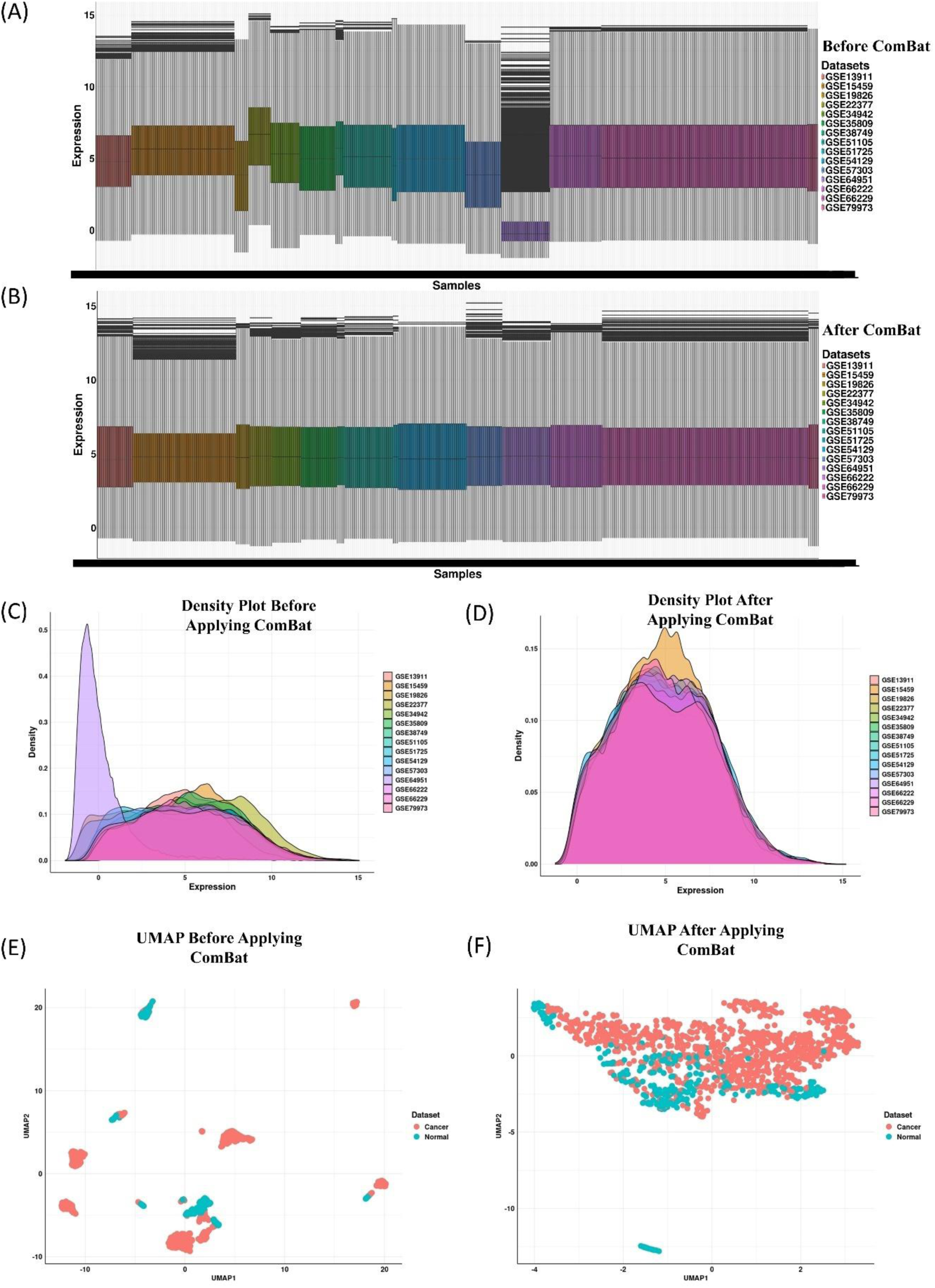
The box plot, density, and UMAP indicates the successful removal of batch effects. Batch-effect elimination is validated through (A) The state before applying ComBat and (B) The state after applying ComBat. Distinct clusters are evident in UMAP representations, showing (C-D) Initial density distribution before and after applying ComBat (E-F) tSNE plot showing the distribution of cancer and normal samples before and after application of ComBat.

### 3.2. Implementation of Machine Learning

Our study utilized common genes identified across fifteen microarray datasets following exclusion criteria. A total of 20,161 genes were obtained, and subsequently, we used the RFE method to select the top 10, 20, 30, 40, and 50 features with the other two feature selections, MI and SelectFromModel tree-based. Since the sample was not balanced, we used SMOTETomek to balance the cancer and normal datasets [43]. Initial samples were 1090 for cancer and 310 for normal samples. After applying SMOTETomek, the samples resulted in an equal number of samples by increasing the samples of the minority class (Cancer =1090 and Normal =1090), as indicated in **Figure 2 (A-B)** and the tSNE plot showing the oversampling of data before and after applying the balancing of dataset using SMOTETomek in **Figure 2 (C)**. We used 3 feature selection methods and 10 machine learning classifiers on the balanced dataset. We tested ten classifiers to classify GC and normal samples (K-Nearest Neighbours (KNN), Decision Tree (DT), XGBoost (XGB), ExtraTrees (ET), Support Vector Classifier (SVC), Gaussian Naive Bayes (GNB), Logistic Regression (LR), Random Forest (RF), AdaBoost (AB), and Multi-layer Perceptron (MLP)). These classifiers were applied to the RFE-selected features (top 10, 20, 30, 40, and 50), MI, and features from tree-based SelectedFromModel. **Figure 2 (D-E)** indicates the accuracy and AUC results as heatmaps for the test dataset. Feature selection RFE_50 (RFE_50 indicates RFE with 50 features) and SVC classifier achieved the highest accuracy and AUC (Accuracy: 97.94 %, AUC = 1), and then RFE_50 and XGB (Accuracy: 97.25 %, AUC = 0.99). However, other combinations also showed up with a better performance. Feature method MI+SVC classifier achieved the highest performance (Accuracy: 97.25 %, AUC = 0.99); similarly for tree-based+AB (Accuracy: 97.02 %, AUC = 0.99) and tree-based+SVC (Accuracy: 97.02 %, AUC = 0.99) achieved the highest performance. Among them, the above-mentioned feature selection technique and the classifier showed the best performance for the complete dataset. Similarly for the cancer driver dataset as shown in figure **Figure 2 (F-G)** indicates the accuracy and AUC results as heatmaps for the test dataset. Feature selection RFE_40 (RFE_40 indicates RFE with 40 features) and RF classifier achieved the highest accuracy and AUC (Accuracy: 96.72 %, AUC = 0.99), and then RFE_40 and SVC (Accuracy: 95.64 %, AUC = 0.98). Feature method MI+SVC classifier achieved the highest performance (Accuracy: 95.87 %, AUC = 0.98); similarly, tree-based+ET (Accuracy: 95.87 %, AUC = 0.99) achieved the highest performance. The ROC curves with AUC values for the test are depicted in **Figure 3 (A-B)** for features selected from RFE, MI, and tree-based relevant features and classifiers. The ROC plot illustrates the classification performance across different decision thresholds, highlighting the model’s ability to distinguish between cancer and normal samples. Each colored curve represents the AUC of a fold from different classifiers, with curves closer to the upper left corner indicating better performance and smaller errors. The blue diagonal dotted line serves as a reference, representing a scenario where the model cannot differentiate between the two classes. In our study, we observed that except for the DT, ET, GNB, and KNN classifiers, all the classifiers had an accuracy >= 90%, and for AUC, it was observed that except for the DT, GNB, and LR classifiers, classifiers had a mean AUC ≥ 0.9 for the complete dataset. A similar pattern was also observed for the cancer driver dataset. **Supplementary Table S2-S7** shows the other performance measures for the complete dataset and cancer driver datasets. The selected features identified by RFE_50, MI, and TB were combined, as depicted in **Figure 3(C)**. In total, 13 features were found to be common in the complete dataset (RFE_50, MI, and TB), while 17 features were common across RFE_40, MI, and TB in the cancer driver dataset as illustrated in **Figure 3(D)**. To assess the performance of these 13 and 17 common genes, models were constructed using the shared genes for both the complete and cancer driver datasets. The models achieved the highest accuracies of 95.8% for the RF classifier and 94.95% for the RF classifier, respectively, using stratified 5-fold cross-validation for complete and cancer driver datasets. Additional evaluation metrics are provided in **Supplementary Table S8-S9**. The RF model attained an AUC value of 0.99 with stratified 5-fold cross-validation for the complete dataset, while the RF model achieved an AUC of 0.98 for the cancer driver dataset, as shown in **Figure 3 (E-F)**. By overlapping the 13 and 17 gene sets, we identified only one common gene, WIF1, as shown in **Supplementary Figure S1 (A)** in the Venn diagram.

**Figure 2:**
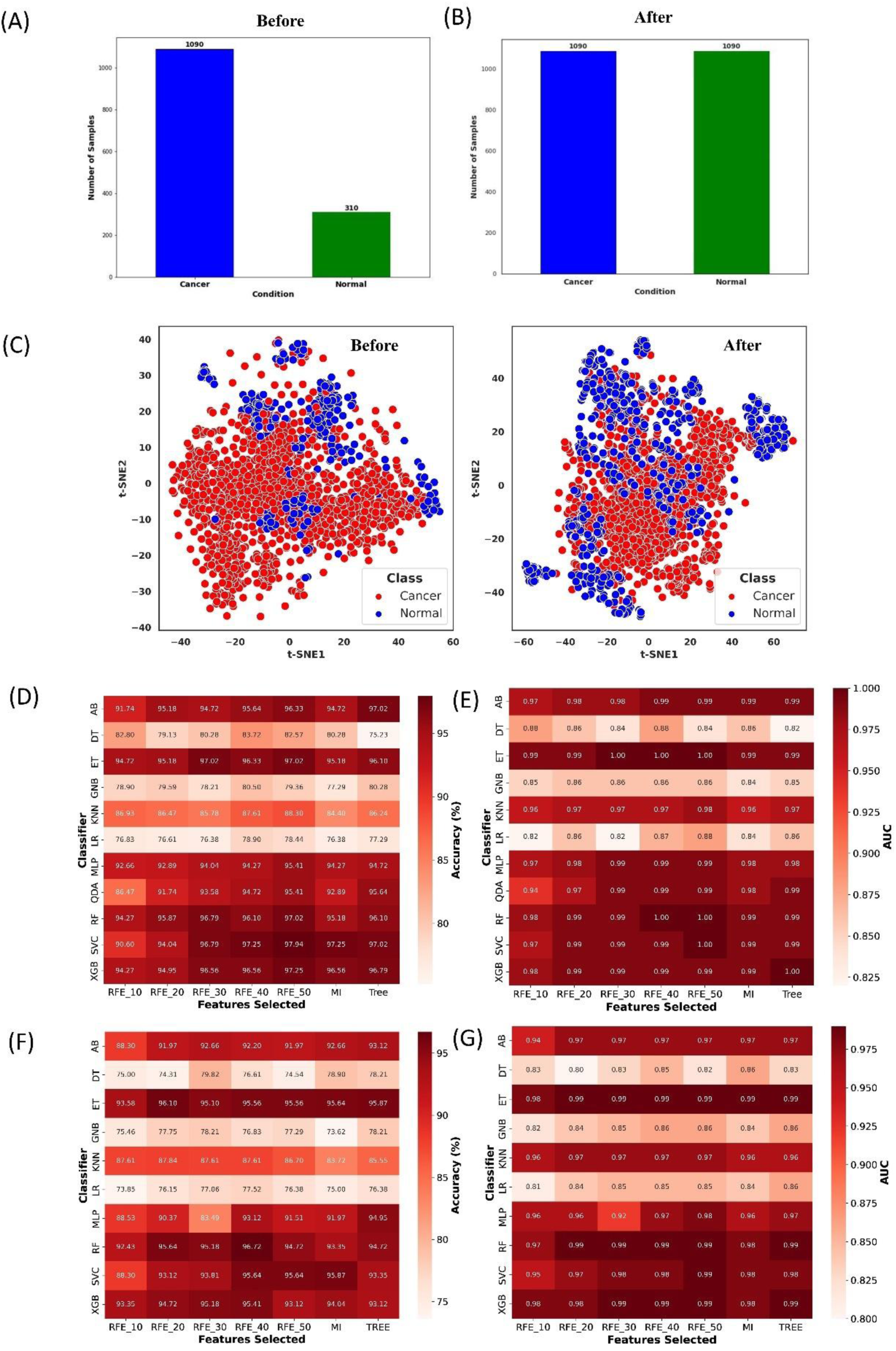
Representative data visualization of cancer driver and complete datasets after employing ML (A-B) Distribution of data before and after applying SMOTETomek link for balancing the dataset for ML (C) tSNE plot of clustering before and after applying SMOTETomek link (D-E) Accuracy and AUC heatmap of the feature selection and classification procedures for complete dataset (F-G) Accuracy and AUC heatmap of the feature selection and classification procedures for cancer driver dataset.

**Figure 3:**
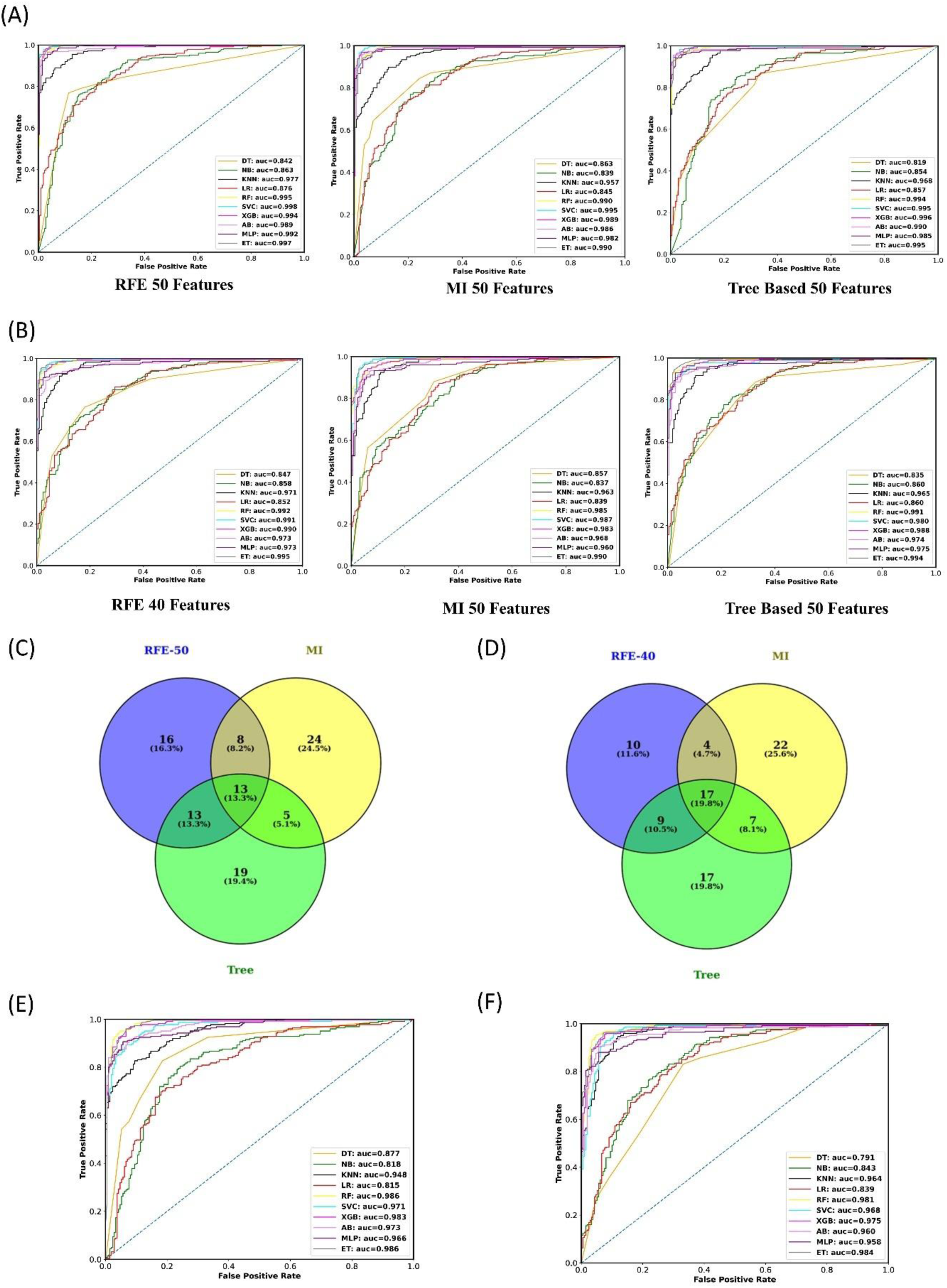
ROC curves were generated for ten machine learning algorithms (A-B) ROC curves using RFE, MI, and tree-based SelectFromModel for the complete and cancer driver dataset. (C) A Venn diagram showing the number of common genes selected from the 50 features from RFE and 50 features from MI and tree-based. (D) A Venn diagram showing the number of common genes selected from the 40 features from RFE and 50 features from MI and tree-based. (E) ROC curve using the 13 common genes from the complete dataset (F) ROC curve using the 17 common genes from the cancer driver dataset.

### 3.3. Identifying Differentially Expressed Genes

In our study, we utilized the limma R package for differential gene expression between cancer and normal gastric samples [44]. We analyzed both driver and complete datasets, and we mapped the genes using features from subset RFE+MI+TB acquired features. Upregulated genes are represented in purple, downregulated genes in green, and non-significant genes in grey (meeting the threshold of logFC ≥ 1 and logFC ≤ −1 and p < 0.05) as indicated in **Figure 4 (A-B)**. When analyzing both driver and complete datasets and mapping them with the differentially expressed (DE) genes, we identified 115 genes with a significant difference in expression levels, defined as having a log fold change (logFC) ≥ 1 or ≤ -1, and a p-value < 0.05. Among these 115 DE genes, 61 were found to be upregulated, while 54 were downregulated. Supplementary Table S8 depicts the DE genes from both driver and complete subsets, along with information on gene regulation, logFC, p-value, pad, and AUC for the 115 genes.

**Figure 4:**
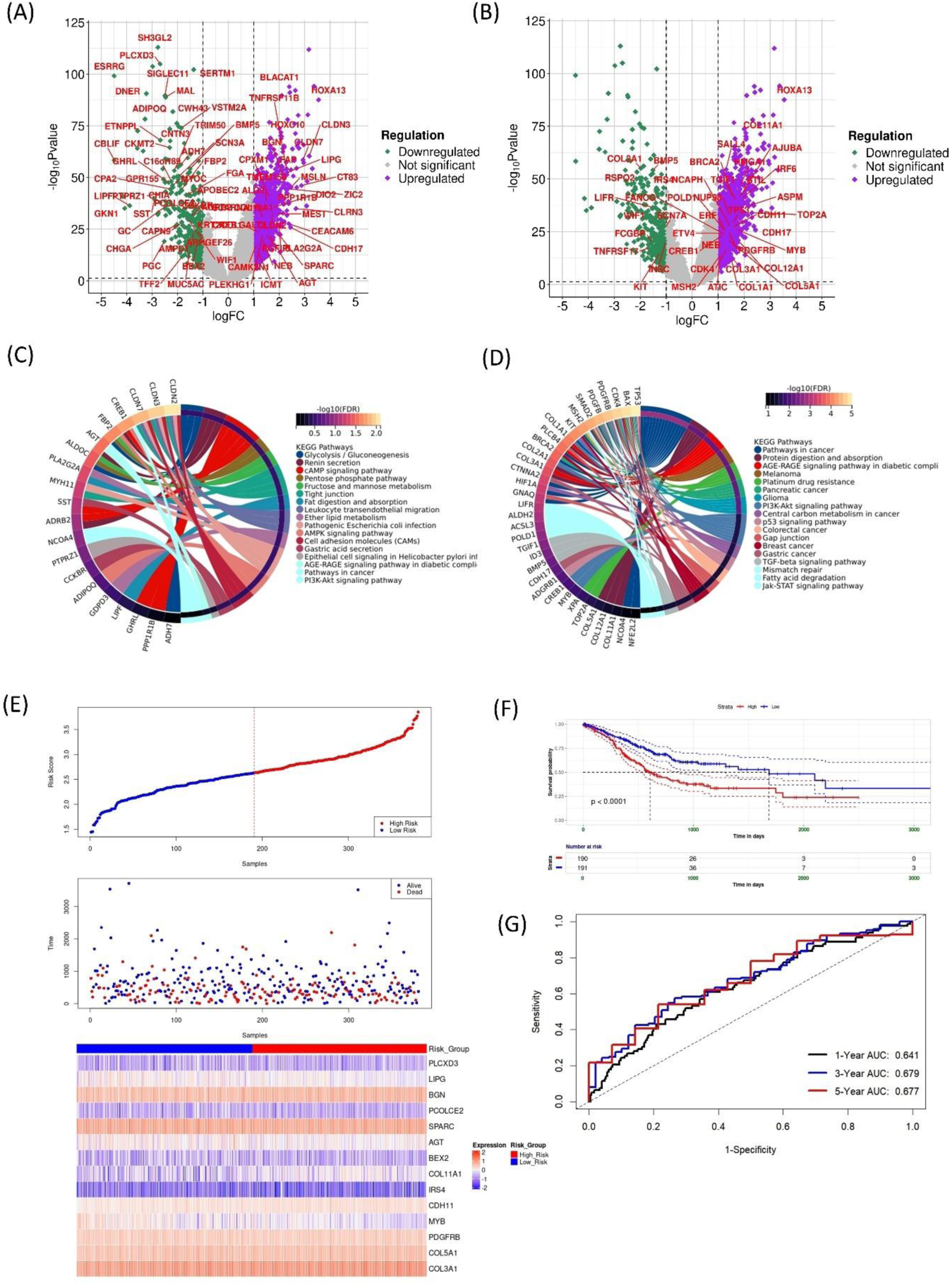
Volcano plots illustrate the differentially expressed genes in gastric cancer. Log fold change is depicted on the axis, with gene transcripts showing a log fold change > 1 marked in purple and those with a log fold change < 1 shown in green. (A) Volcano plot of genes selected from the complete dataset and (B) Volcano plot of genes selected from the cancer driver dataset. (C) Chord plot to visualize the relationships between genes and their associated pathways within genes from the complete dataset and (D) cancer driver genes datasets. (E) The patients diagnosed with gastric cancer (TCGA-STAD) were stratified into high- and low-risk groups. Their respective risk scores, survival outcomes, and the expression levels of the fourteen genes are depicted. (F) Patients with gastric cancer classified in the high-risk category exhibited poorer survival rates. (G) ROC curve of the risk score model for the TCGA-STAD dataset.

Additionally, we conducted pathway analysis for both complete and driver datasets, identifying key pathways. In the complete datasets, we observed significant pathways such as glycolysis/gluconeogenesis, renin secretion cAMP signaling pathway, pentose phosphate pathway, and followed other pathways as shown in **Figure 4 (C)**. Cancer driver, datasets exhibited top pathways, including pathways in cancer, protein digestion and absorption, AGE-RAGE signaling pathway, and other pathways indicated in **Figure 4 (D)**.

### 3.4. Survival Analysis

We conducted a prognostic model to calculate the risk score of each patient using the TCGA-STAD dataset and two test datasets (GSE34942 and GSE15459). A lasso penalized Cox regression was applied to 115 differentially expressed genes, identified after overlapping DEGs from the complete and the cancer driver datasets. This analysis narrowed the selection down to fourteen significant genes, as shown in **Supplementary Figure S1(B-C).** Lasso-penalized Cox regression combines variable selection and estimation in one method. It uses the Cox proportional hazards model, a popular tool for analyzing time-to-event data, along with a lasso penalty to shrink less important features’ coefficients towards zero, potentially eliminating some entirely. This aids in identifying relevant features and reducing model complexity.

Patients were then stratified into high and low-risk groups based on the median risk score cut-off value. The risk score was calculated using coefficients from the model and gene expression values, with each gene’s expression multiplied and summed for the fourteen selected genes, as shown in **Figure 4 (E)**. Subsequently, we used Kaplan-Meier curves to evaluate the predictive ability of the risk score model and assess survival outcomes. The results indicated that the prognosis of patients with a high-risk score had poorer prognosis using the fourteen genes (p < 0.0001). Notably, the high-risk group in the samples of the TCGA-STAD dataset exhibited significantly poorer overall survival compared to the low-risk group, as illustrated in **Figure 4 (F)**. The 14 genes identified through stepwise multivariate regression analysis were used to construct a prognostic gene signature model, as depicted in **Supplementary Figure S1(D).** The area under the curve (AUC) for the ROC analysis at 1, 3, and 5 years was 0.64, 0.679, and 0.67, respectively, for TCGA-STAD, as shown in **Figure 4(G)**, indicating good predictive capability of the model. This prognostic model was further validated using the GSE34942 and GSE15459 datasets. **Supplementary Figures S2(A-B)** display the risk score distribution, gene expression values, and survival status of patients. Similarly, the results indicated that patients in the high-risk score group had a lower survival rate, as shown in **Supplementary Figures S2(C-D).** The AUC for the ROC analysis at 1, 3, and 5 years was 0.63, 0.83, and 0.82 for the GSE34942 dataset, and 0.69, 0.74, and 0.76 for the GSE15459 dataset, as shown in **Supplementary Figures S2(E-F).** These findings collectively suggest that the established prognostic model has good predictive performance.

### 3.5. Expression analysis of Fourteen potential Signatures

To assess the expression patterns and clustering performance of fourteen genes, we compared their expression levels between GC and normal samples. In addition to normal samples, we also included normal-GTEx samples in our analysis. We visualized the clustering performance using a heatmap and PCA plot based on the expression signatures of the fourteen genes **Figure 5 (A-B)**. In the heatmap, samples were represented on the horizontal axis and genes on the vertical axis. Out of 14 genes, 10 genes were upregulated (LIPG, BGN, SPARC, AGT, COL11A1, CDH11, MYB, PDGFRB, COL5A1, COL3A1), and 4 genes were downregulated (PLCXD3, PCOLCE2, BEX2, IRS4). Distinct clusters were observed for GC and normal samples in both the heatmap and PCA plot. We considered subgroup information during the clustering analysis. **Figure 5 (B)** shows a slight overlap between normal samples and GTEx samples, but both sets of normal samples could be completely distinguished from tumor samples. Specifically, all fourteen of these genes exhibited strong discriminatory power in distinguishing between tumors and normal samples, with an area under the curve (AUC) ranging from 0.60 to 0.80 **Figure 5 (C)**. The AUC value is higher than 0.60 for all fourteen genes, suggesting that they are promising indicators for predicting tumor prognosis in GC. Through a feature selection approach in GC, these fourteen genes were identified as promising classifiers between cancerous and normal samples. To examine the expression patterns of these genes in GC, normal samples, and GTEx normal samples, differential expression was analyzed using boxplots in **Figure 5 (D)**, revealing a significant difference between cancer and normal samples (p < 0.05).

**Figure 5:**
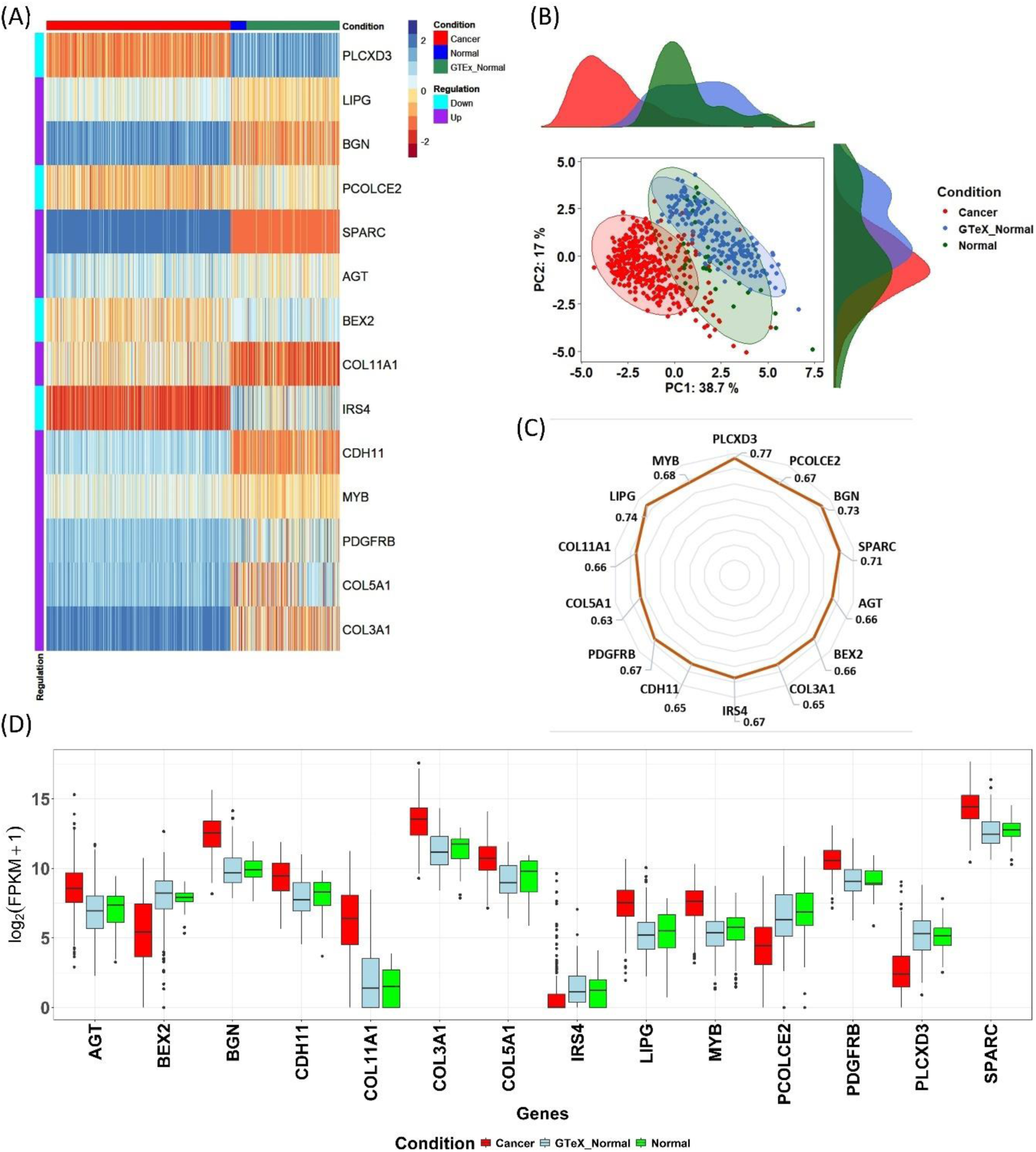
(A) A heatmap indicating the expression patterns across cancer, normal, and GTeX normal samples, accompanied by identifying up- and down-regulated genes for fourteen specific genes. (B) PCA plot depicting the distribution of cancer, normal, and GTeX normal samples. (C) Evaluation of the area under the curve (AUC) values for all fourteen genes, demonstrating that each gene possesses an AUC greater than 0.60. (D) Comparative analysis through boxplots illustrating the expression levels of the fourteen genes across cancer, normal, and GTeX normal samples.

After analysing the differential expression of genes, all fourteen mentioned genes were investigated individually to assess their association with overall survival. The Kaplan-Meier survival curves indicated that elevated expression levels of PLCXD3, PCOLCE2, BGN, SPARC, AGT, BEX2, COL3A1, IRS4, CDH11, PDGFRB, COL5A1, COL11A1 were associated with a decreased likelihood of overall survival, whereas higher expression levels of MYB and LIPG were linked to improved survival outcomes. Kaplan-Meier analysis was performed to identify genes with significant differences in expression levels that could effectively distinguish between patients based on their survival outcomes in GC, as depicted in **Figure 6 (A-N)**.

**Figure 6:**
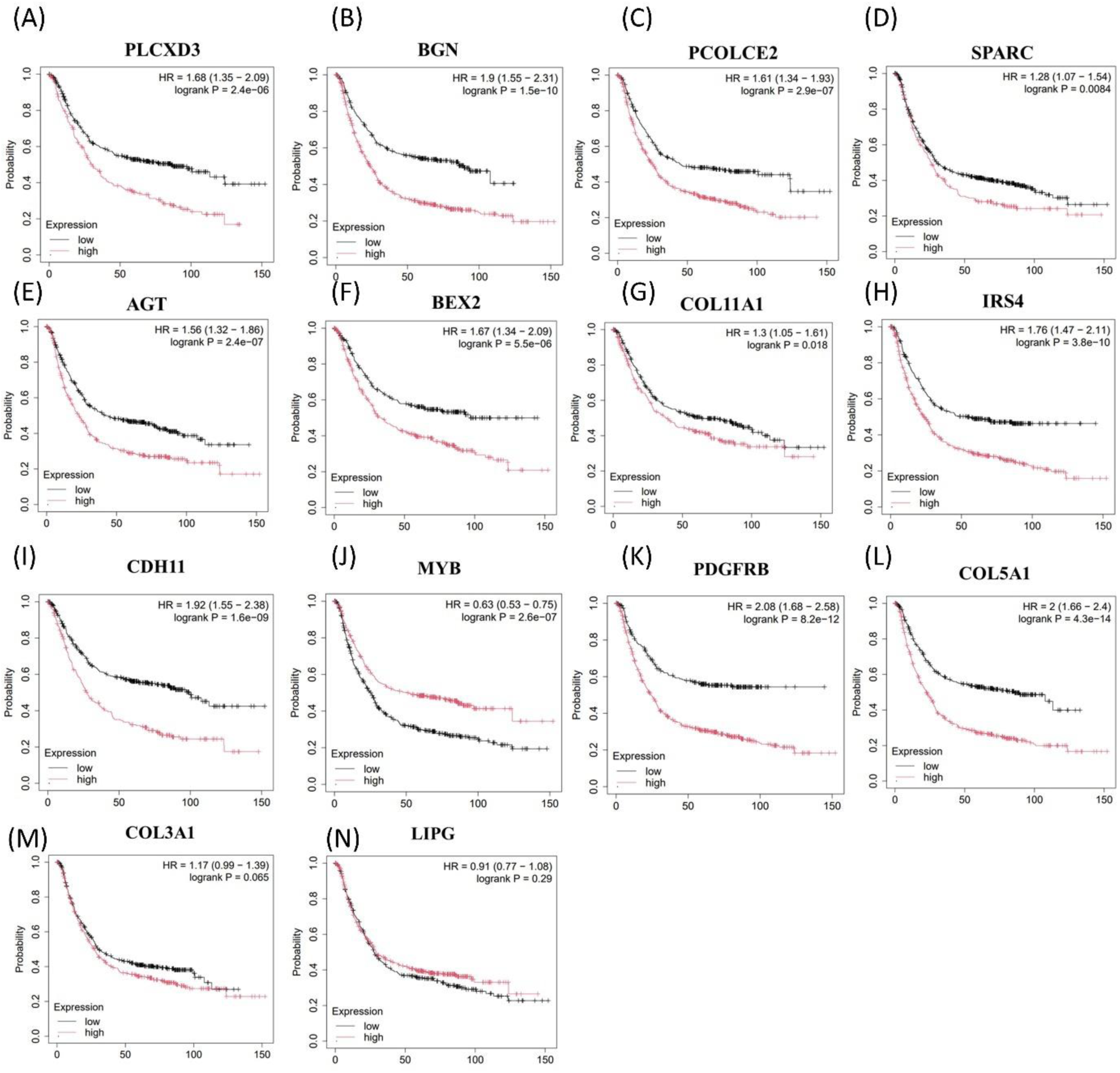
(A-N) Kaplan-Meier plot depicting overall survival for the fourteen genes.

### 3.6. Co-expression network analysis using WGCNA

Co-expressed genes often participate in related signaling pathways and exhibit similar biological functions. To identify co-expressed genes among the 115 identified genes, a co-expression network analysis was conducted using WGCNA (Weighted Gene Co-expression Network Analysis) on microarray data from all 15 datasets combined. Fisher’s exact test was used to identify significant modules containing the 115 genes. The analysis began by transforming the expression matrix into a topological overlap matrix using a soft-thresholding power of β = 6, as shown in **Figure 7(A-C)**. Genes were then grouped into different modules through dynamic tree cutting. Subsequently, an association analysis between clinical traits and gene modules was performed. Using Fisher’s exact test, three significant modules were identified, containing 14 key genes (PLCXD3, PCOLCE2, BGN, SPARC, AGT, BEX2, COL3A1, IRS4, CDH11, PDGFRB, COL5A1, COL11A1, LIPG, MYB) from previous analysis. These genes showed significant associations within the three modules. Specifically, the turquoise module demonstrated a positive correlation with cancer traits (cor = 0.17, p = 2E-09), the blue module showed a negative correlation (cor = -0.17, p = 2E-10), and the red module exhibited a weaker correlation with cancer traits as shown in **Figure 6(D)**. Genes from the blue, red, and turquoise modules were selected based on specific criteria. To further explore the functions of these key genes, KEGG pathway enrichment analysis was conducted for each module **Figure 7(E-G)**. The most significantly enriched pathways in the blue module were DNA replication, mismatch repair, homologous recombination, and cell cycle. For the red module, the top pathways were propanoate metabolism, valine, leucine, and isoleucine degradation, oxidative phosphorylation, and ECM-receptor interaction. In the turquoise module, the enriched pathways included dilated cardiomyopathy, hypertrophic cardiomyopathy, axon guidance, and focal adhesion.

**Figure 7:**
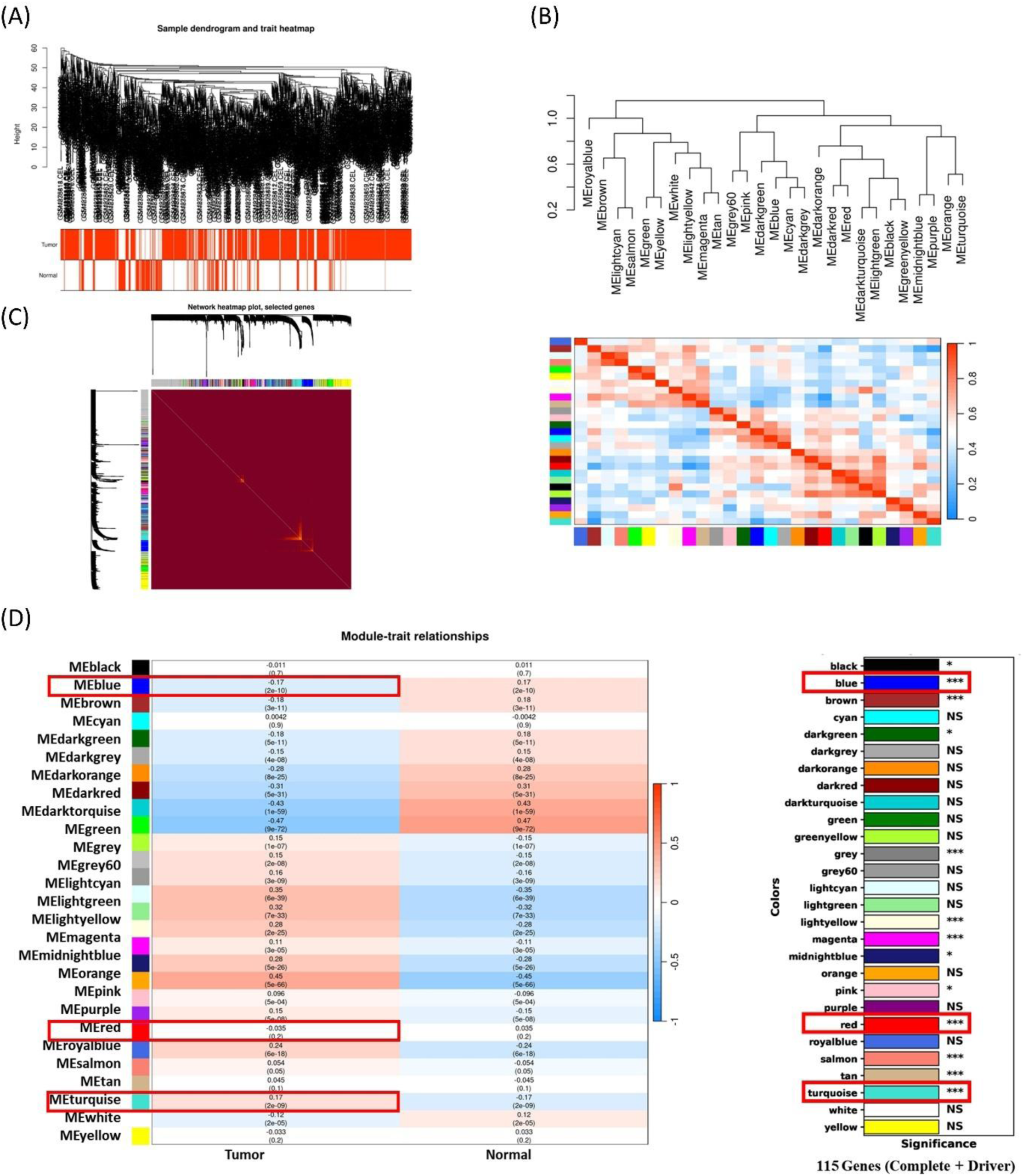
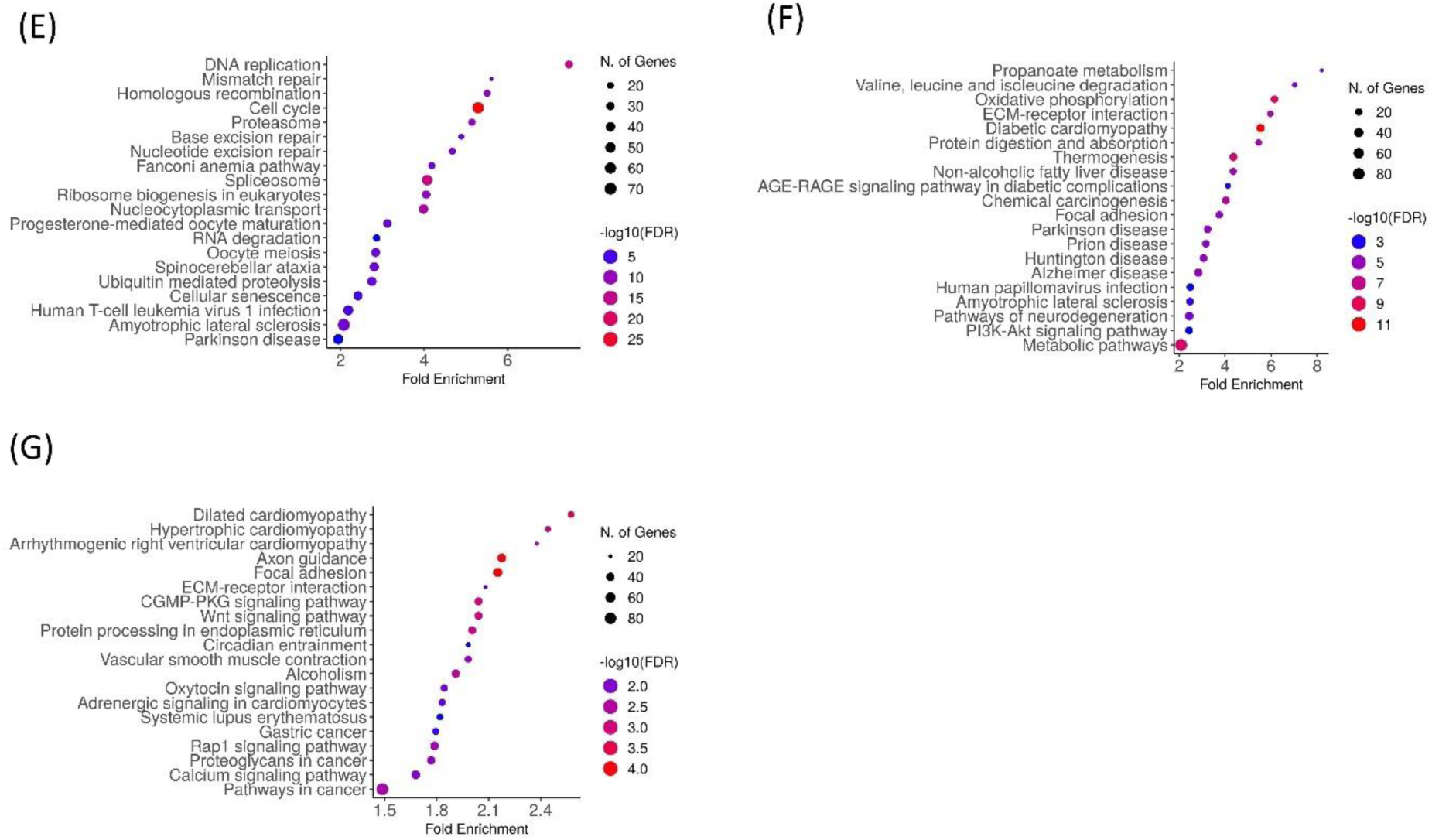
(A) The clustering dendrogram illustrates the hierarchical clustering of samples based on their Euclidean distance. (B) A heatmap plot visualizes the gene network structure. (C) The Topological Overlap Matrix (TOM) plot demonstrates that the modules within the network are independent of each other, indicating that gene expression within each module is also relatively independent. (D) This heatmap displays the correlation between gene modules and clinical traits, with p < 0.05 indicating statistical significance. Significant modules containing key prognostic genes are highlighted. (E-G) KEGG pathway enrichment analysis of blue, red, and turquoise modules.

## 4. Discussion

In the realm of biology and life sciences, the development of high-throughput techniques and Next Generation Sequencing (NGS) analysis has led to the generation of numerous omics datasets, particularly in cancer research. Feature selection methods have become crucial in identifying important subsets of features from high-dimensional datasets, which is essential for advancing precision medicine, including cancer diagnosis and treatment [45–46]. In our investigation, we employed both Recursive Feature Elimination (RFE) method for feature selection and identified fourteen genes with promising performance in distinguishing GC from normal colon tissues. Further, expression analysis was conducted based on a combined dataset from both the TCGA-STAD cohort and GTEx normal stomach samples [47–48].

Traditionally, identifying differentially expressed or abundant molecules such as genes, proteins, or metabolites between tumor and control groups often yields hundreds or even thousands of candidates. However, identifying biomarkers from this large pool can be challenging. To address this issue, additional methods are needed to efficiently select key gene signatures that accurately classify the disease compared to normal samples [49–50].

GC is a highly aggressive disease often diagnosed in advanced stages, presenting a significant global health challenge despite advancements in treatment modalities [51]. Unfortunately, the prognosis for GC remains dismal, underscoring the necessity for further investigation to identify prognostic biomarkers and potential therapeutic targets. In our study, we utilized a combination of traditional approaches and machine learning techniques. Initially, we identified differentially expressed genes, while simultaneously employing machine learning algorithms to identify the most relevant subset for better classification accuracy. We used the three feature selection methods (RFE, MI and TB) to identify the best features to classify the cancer and normal samples for the complete and cancer driver gene datasets. After obtaining the feature from the individual feature selection for both the complete and cancer driver gene datasets, we identified 13 and 17 feature genes that were common for the 3 feature selection methods. We performed the AUCROC curve to evaluate the diagnostic potential of these genes, and all the genes showed an AUC greater than 0.65. Further, we found WIF1 as the only common gene from the 13 and 17 genes obtained from complete and cancer driver gene datasets. Simultaneously, we also mapped the genes obtained from both the dataset and the genes obtained from DEG analysis from limma. We obtained 115 genes that were dysregulated with logFC > 1 and logFC < -1 with a value < 0.05 threshold. With these 115 genes, we further used the lasso penalized Cox regression model to obtain the prognostic gene signatures. Ultimately, we identified a set of 14 genes that exhibited powerful discrimination between GC and normal tissues. Importantly, we validated these findings independently using two GC microarray datasets, confirming both the expression alterations and the classification performance of the selected gene sets. Furthermore, survival analysis highlighted PLCXD3, PCOLCE2, BGN, SPARC, AGT, BEX2, COL3A1, IRS4, CDH11, PDGFRB, COL5A1, COL11A1, LIPG, and MYB as prognostic genes significantly associated with overall survival in GC patients.

When ranking the genes, PLCXD3 displayed the highest AUC value with 0.77 among the 14 genes. We further did co-expression gene analysis using WGCNA as co-expressed genes often participate in common pathways, and 3 modules were found to be significant in which these 14 genes were present (blue, red, and turquoise). However, the red module didn’t show up with a significant module trait relationship and correlation with the cancer trait. The turquoise module genes were involved in pathways such as dilated cardiomyopathy, hypertrophic cardiomyopathy, axon guidance, and focal adhesion, and in the blue module, DNA replication, mismatch repair, homologous recombination, and cell cycle.

PLCXD3 encodes a phospholipase that hydrolyzes phospholipids into fatty acids. While the precise role of PLCXD3 in cancer remains unclear, it has been identified as a protective factor in lung cancer, with its expression showing a negative correlation with cancer risk [52]. Notably, in our study, PLCXD3 demonstrated the highest AUC among the 14 genes analyzed. SPARC encodes a member of the extracellular matrix glycoprotein family that binds calcium ions. This protein has been shown to facilitate tumor cell detachment, invasion, metastasis, basement membrane degradation, endothelial cell migration, angiogenesis, cell growth stimulation, and alterations in matrix composition [53]. A study also revealed that both SPARC protein and mRNA expression levels are elevated in GC [54]. In our study, we observed the expression of SPARC as an upregulated gene. PCOLCE2, a collagen-binding protein, was found to be downregulated in our study [55]. Another study utilizing whole exome sequencing identified mutations in PCOLCE2 among family members with GC [56]. These findings suggest a potential role for PCOLCE2 in GC, but further research is necessary to elucidate its precise function and impact on the disease. BGN, also known as Biglycan, has been found to be upregulated in various cancers, including pancreatic, gastric, prostate, and colon cancers. Its overexpression is linked to aggressive tumor growth and metastasis [57]. LIPG, also known as Endothelial Lipase, was found to be upregulated in our study. A previous study on GC identified LIPG as a potential urinary biomarker, demonstrating high sensitivity and specificity for the disease [58]. Insulin receptor substrate 4 (IRS4) was found to be downregulated in our study. A recent study demonstrated that hsa_circ_0023409 activates the IRS4/PI3K/AKT pathway by acting as a sponge for miR-542-3p, thereby promoting the development and progression of GC [59]. CDH11, a type II cadherin, has been found to be significantly overexpressed in GC tissues compared to normal gastric tissues in a previous study. Similarly, our study also observed higher CDH11 expression in GC. Additionally, that study reported that CDH11 expression was elevated in late-stage compared to early-stage GC, with increased levels of CDH11 being associated with tumor progression and poor prognosis in patients with GC [60]. BEX2 (Brain expressed X-linked 2) was found to be downregulated in our study, and a recent study also identified BEX2 as a poor prognostic factor in GC cohorts [61]. MYB family members are often aberrantly expressed in human cancers and play crucial roles in growth, differentiation, and survival. Overexpression of MYB has been observed in acute myeloid leukemia, non-Hodgkin lymphoma, colorectal cancer, and breast cancer [62–65]. A study also revealed MYB overexpression in GC, which is negatively regulated with the prognosis [66]. PDGFR-β, expressed in stromal and perivascular cells, plays a role in angiogenesis, tumor proliferation, and is considered a cancer prognostic marker [67–68]. In colorectal liver metastasis, Increased expression of PDGFR-β promotes epithelial-to-mesenchymal transition, enhancing tumor aggressiveness [69]. In GC, high PDGFR-β levels correlate with increased M2 macrophage infiltration and negatively impact prognosis by promoting angiogenesis and modulating the tumor immune microenvironment [69]. COL5A1 has been reported to promote proliferation and metastasis of multiple tumors, such as lung cancer, GC, breast cancer, and renal cell carcinoma [70–73]. COL5A1 Promotes the progression of GC by affecting the immune infilteration as acting as a ceRNA and reversely sponge miR-137-3p to upregulate the expression of FSTL1 [74]. Another member of collagen family COL3A1(Collagen Type III Alpha 1 Chain) has shown the malignant progression and drug resistance in a variety of cancers as it plays vital role in cell adhesion, migration, proliferation, and differentiation via interactions with cell-surface receptor integrins [75]. In GC, the upregulation of COL3A1 is closely associated with poor prognosis. Additionally, COL3A1 has been recognized as a promising independent predictor for the prognosis of head and neck cancer, as well as colorectal cancer [76–78]. An additional member of the collagen family Collagen 11 alpha 1 chain (COL11A1) is known to regulate tumor progression in various cancers, including esophagus, colorectal, gastric, glioma, lung, ovarian, pancreatic, salivary gland, head and neck, and renal cancers [79]. An in-silico analysis revealed that COL11A1 and COL1A1 genes are potenial diagnostic biomarkers in Breast, colorectal and GCs [80]. A study revealed upregulation of COL11A1 offered poor prognosis and increased cell proliferation, migration, and invasion and was also associated with the upregulation of proliferation-related genes, including CDK6, TIAM1, ITGB8, and WNT5A [81]. AGT is a component of Renin-Angiotensin system and a signature gene of lipid metabolism and cell death. Enhanced AGT expression association with higher levels of T cells CD4 memory resting and lower activated B cells which suggests AGT potentially influences the immune microenvironment of COAD patients [82]. Upregulation of AGT was also observed in GC samples than the normal samples in TCGA dataset and results also showed that upregulated AGT expression was significantly correlated with poor OS [83].

Understanding the precise functions of these genes and how their changes contribute to cancer progression allows researchers to study cancer in a more focused manner. This enhanced understanding can then facilitate the creation of personalized treatments customized to the individual genetic characteristics of each patient’s cancer, thereby enhancing the effectiveness and precision of treatment approaches.

In summary, using a combination of feature selection techniques has demonstrated strong reliability and effectiveness in identifying accurate and biologically relevant biomarker genes. The consistency between the results of our approach, in combination with bioinformatics downstream analysis, and previous studies reinforces the validity of our approach. This supports the use of different feature selection methods together to identify signature genes, rather than depending on a single method.

## 5. Conclusion

Our study utilized the 3 feature selection methods from filter-based and wrapper methods to identify a signature of fourteen genes that effectively distinguish GC from normal samples. Among the genes analyzed, ten exhibited significant upregulation in cancer samples, while four showed downregulation. Survival analysis indicated that patients with high-risk profiles had poorer prognoses, particularly associated with these fourteen genes. Our study emphasizes the importance of employing feature selection techniques, especially when integrating diverse datasets from microarray experiments, to advance precision medicine in cancer research. We expect that unraveling the roles of these prognostic genes in GC will provide valuable insights into potential targets for future interventions and therapies. However, a limitation of these identified biomarkers is their origin from tissue samples, necessitating invasive extraction methods. Additional validation is necessary, and a similar approach can be applied to blood samples, which require non-invasive methods for reliable analysis.

## Author contributions

AS, PKS, and RV collaboratively designed and conceptualized the study. RV performed the analysis and prepared the initial manuscript draft. AS and PKS authorized further analyses, contributed to editing, and reviewed the manuscript. All authors provided final approval for the manuscript, collectively contributing to the research article and endorsing the submitted version.

## Data Availability

The datasets utilized in our investigation have been delineated within the study and can be accessed via the accession codes GSE66229, GSE54129, GSE13911, GSE19826, GSE79973, GSE51725, GSE15459, GSE51105, GSE35809, GSE57303, GSE34942, GSE22377, GSE38749, GSE66222, GSE64951 from the NCBI GEO dataset repository. Additionally, TCGA-STAD expression data can be obtained from the GDC portal.

## Abbreviations

STAD: Stomach adenocarcinoma
NGS: Next-generation sequencing
ML: Machine Learning
DEGs: Differentially expressed genes
SVC: Support vector classifier
kNN: K-nearest neighbor
ET: ExtraTrees
DT: Decision tree
MLP: Multi-layer Perceptron
RF: Random forest
LR: Logistic Regression
GNB: Gaussian Naive Bayes
AB: AdaBoost
XGBoost: eXtreme gradient boosting
OS: Overall survival
GEO: Gene Expression Omnibus
TCGA: The cancer genome atlas
MI: Mutual Information
RFE: Recursive Feature Elimination
KM: Kaplan-Meier plot
GTEx: The Genotype-Tissue Expression

## Acknowledgments

The authors extend their gratitude to the Shiv Nadar Institution of Eminence (SNIoE) for offering the essential research framework and infrastructure that made this study possible.

