## Supplementary File for "Discovery of Prognostic Biomarkers in Gastric Cancer Through Machine Learning and Bioinformatics Analysis of Gene Expression Data"

***Supplementary Methodology:***

### 1.1 Feature Selection

We utilized the SMOTETomek function in Python to up-sample the minority class since the dataset was highly imbalanced (310 for Normal and 1090 for Cancer). This imbalanced dataset might give a biased result. Therefore, we applied SMOTETomek before applying the different feature selection approach as it provided an equal set of normal and cancer samples. After that, we for constant features present in the dataset, followed by removing quasi-constant features from the dataset, which ultimately led to 1168 features for cancer-driver datasets and 19769 to complete datasets. Further, we used Pearson's correlation coefficient algorithm to reduce redundant features in our dataset (For the complete and driver datasets, genes with a correlation coefficient of $\geq 0.90$ were excluded), which resulted in removing 75 correlated features for the complete dataset and 1 feature for cancer driver genes. So, it finally provided 19,694 features for complete datasets and 1167 features for cancer-driver datasets. We also employed the Area under the Receiver Operating Characteristic (AUC) based feature selection technique, wherein we computed the AUC for each individual feature (AUC value $\geq 0.60$ selected for the cancer-driver dataset and complete dataset). This narrowed the features in the complete dataset to 5311 features and 349 features in cancer driver datasets. Finally, Recursive Feature Elimination (RFE) was employed on both datasets to extract the top 10, 20, 30, 40, and 50 gene features and the top 50 features from both mutual information (MI) and tree-based (TB) feature selection.

RFE is a type of wrapper method suitable for regression and classification problems, and it internally employs filter-based feature selection. It begins by searching all features in the training dataset and removing the irrelevant features until the desired features are left. Mutual Information (MI) is a filter method used in feature selection algorithms. It simultaneously evaluates the relevance of features to the target classes and assesses the redundancy between features. MI can also determine the association between a random feature and another feature. The tree-based feature selection method using random forest is also a wrapper method. This approach combines feature selection with model training. These methods uses the intrinsic properties of decision trees to assess feature importance while concurrently constructing the predictive model.

Subsequently, we applied ten different machine learning classifiers such as (1) K-Nearest Neighbours (KNN), (2) Decision Tree (DT), (3) XGBoost (XGB), (4) ExtraTrees (ET), (5) Support Vector Classifier (SVC), (6) Gaussian Naive Bayes (GNB), (7) Logistic Regression (LR), (8) Random Forest (RF), (9) AdaBoost (AB), and (10) Multi-layer Perceptron (MLP) to classify gastric cancer and normal samples on each feature subset obtained from RFE, MI and TB. The Complete and Driver datasets underwent a random split, with an 80:20 ratio where 80% of the data was used for the training machine learning models and the remaining 20% for testing purposes. Every algorithm came with a specific set of hyperparameters that needed to be adjusted for each dataset. We optimized these hyperparameters through grid search combined with 5-fold stratified cross-validation on a portion of the training data. Following this, the refined model was tested on separate holdout data to assess and compare the performance of the five algorithms. The prediction accuracy achieved during cross-validation was averaged to establish the final accuracy. Furthermore, we evaluated various performance metrics on the test datasets.

#### 1.2 Performance Measures

Evaluation metrics play a major role in assessing the effectiveness of machine learning models, and it's important to select metrics that align with our objectives. In our analysis of classification models, we employed sensitivity, specificity, accuracy, Matthew's correlation coefficient (MCC), receiver operating characteristic (ROC) curve, area under the curve (AUC), recall, and precision. Among these metrics, ROC/AUC, MCC, sensitivity, and precision offer valuable insights into the model's performance. We utilized standard equations to calculate these metrics, which are commonly employed in evaluating classification models. These equations allow us to quantify the model's performance in terms of sensitivity, specificity, accuracy, and MCC, providing a comprehensive understanding of its effectiveness.

$$(Sensitivity) = \frac{TP}{TP + TN} * 100$$

$$(\text{Specificity}) = \frac{TN}{TN + FP} * 100$$

$$(\text{Accuracy}) = \frac{(TP + TN)}{(TP + FP + TN + FN)} * 100$$

$$(\text{AccuracyMatthew's correlation coefficient (MCC)}) = \frac{(TP * TN) - (FP * FN)}{\sqrt{(TP + FP)(TP + FN)(TN + FP)(TN + FN)}} * 100$$

Here, False positive, False negative, True positive, and True negative predictions are denoted by FP, FN, TP, and TN, respectively.

#### 1.3 Gene Expression and Statistical Analysis

To examine the expression profiles of the eight reduced genes in both cancer and normal tissue samples, we also incorporated GTEx normal samples specific to gastric tissue. The Genotype-Tissue Expression (GTEx) project is an extensive research endeavor focused on understanding human gene expression and its regulation across a range of tissues. The study provides a comprehensive overview of gene expression patterns and investigates the impact of genetic variations on gene regulation in various human tissues. Initially, gene expression levels were measured in FPKM units (fragments per kilobase of transcript per million mapped reads). Due to significant variability in FPKM values, a logarithmic transformation was applied by adding a constant value of 1 to each FPKM measurement. Furthermore, the data underwent z-score normalization using the caret package in R to prepare it for further analysis. Heatmap visualization was then employed to depict the expression patterns of eight genes, highlighting those that were either upregulated or downregulated. Additionally, a PCA plot was generated to identify trends across cancerous, normal, and GTEx normal samples. Predictive values for each gene were assessed using AUC calculations. The gene expression patterns were further depicted via boxplots, with statistical significance set at $p$ value $< 0.05$ using paired t-tests to assess differences between cancer and normal samples.

***Supplementary Tables***

**Table S1.** The NCBI-GEO Samples were used for this study.

| Series | Tumor | Normal | Platform ID |
|---|---|---|---|
| GSE66229 | 300 | 100 | GPL570 |
| GSE54129 | 111 | 21 | GPL570 |
| GSE13911 | 38 | 31 | GPL570 |
| GSE19826 | 12 | 15 | GPL570 |
| GSE79973 | 10 | 10 | GPL570 |
| GSE51725 | 8 | 2 | GPL570 |
| GSE15459 | 200 | 0 | GPL570 |
| GSE51105 | 94 | 0 | GPL570 |
| GSE35809 | 70 | 0 | GPL570 |
| GSE57303 | 70 | 0 | GPL570 |
| GSE34942 | 56 | 0 | GPL570 |
| GSE22377 | 43 | 0 | GPL570 |
| GSE38749 | 15 | 0 | GPL570 |
| GSE66222 | 0 | 100 | GPL570 |
| GSE64951 | 63 | 31 | GPL570 |

**Table S2.** The performance of gastric cancer classification was developed using 50 features where sensitivity, specificity, and accuracy were measured in percentage using test datasets from complete datasets using RFE.

| Classifier | Sensitivity | Specificity | Accuracy | AUC | MCC |
|---|---|---|---|---|---|
| AB | 95.11 | 97.63 | 96.33 | 0.99 | 0.93 |
| DT | 76.89 | 88.63 | 82.57 | 0.84 | 0.66 |
| ET | 97.33 | 96.68 | 97.02 | 1 | 0.94 |
| GNB | 84.89 | 73.46 | 79.36 | 0.86 | 0.59 |
| KNN | 78.22 | 99.05 | 88.3 | 0.98 | 0.79 |
| LR | 82.22 | 74.41 | 78.44 | 0.88 | 0.57 |
| MLP | 93.33 | 97.63 | 95.41 | 0.99 | 0.91 |
| RF | 96.89 | 97.16 | 97.02 | 1 | 0.94 |
| SVC | 97.78 | 98.1 | 97.94 | 1 | 0.94 |
| XGB | 97.33 | 97.16 | 97.25 | 0.99 | 0.94 |

**Table S3**. The performance of gastric cancer classification was developed using 50 features where sensitivity, specificity, and accuracy were measured in percentage using test datasets from complete datasets using MI.

| Classifier | Sensitivity | Specificity | Accuracy | AUC | MCC |
|---|---|---|---|---|---|
| AB | 92.44 | 97.16 | 94.72 | 0.99 | 0.9 |
| DT | 84.44 | 75.83 | 80.28 | 0.86 | 0.61 |
| ET | 95.11 | 95.26 | 95.18 | 0.99 | 0.9 |
| GNB | 82.67 | 71.56 | 77.29 | 0.84 | 0.55 |
| KNN | 75.11 | 94.31 | 84.4 | 0.96 | 0.7 |
| LR | 78.67 | 73.93 | 76.38 | 0.84 | 0.53 |
| MLP | 92 | 96.68 | 94.27 | 0.98 | 0.89 |
| RF | 94.67 | 95.73 | 95.18 | 0.99 | 0.9 |
| SVC | 96.89 | 97.63 | 97.25 | 0.99 | 0.93 |
| XGB | 95.11 | 98.1 | 96.56 | 0.99 | 0.93 |

**Table S4.** The performance of gastric cancer classification was developed using 50 features where sensitivity, specificity, and accuracy were measured in percentage using test datasets from complete datasets using SelectFromModel (Tree-based).

| Classifier | Sensitivity | Specificity | Accuracy | AUC | MCC |
|---|---|---|---|---|---|
| AB | 95.11 | 99.05 | 97.02 | 0.99 | 0.94 |
| DT | 81.33 | 68.72 | 75.23 | 0.82 | 0.51 |
| ET | 96.89 | 95.26 | 96.1 | 0.99 | 0.92 |
| GNB | 85.33 | 74.88 | 80.28 | 0.85 | 0.61 |
| KNN | 75.11 | 98.1 | 86.24 | 0.97 | 0.75 |
| LR | 79.56 | 74.88 | 77.29 | 0.86 | 0.55 |
| MLP | 91.56 | 98.1 | 94.72 | 0.98 | 0.9 |
| RF | 96.44 | 95.73 | 96.1 | 0.99 | 0.92 |
| SVC | 96.44 | 97.63 | 97.02 | 0.99 | 0.93 |
| XGB | 96.44 | 97.16 | 96.79 | 1 | 0.94 |

**Table S5**. The performance of gastric cancer classification was developed using 50 features where sensitivity, specificity, and accuracy were measured in percentage using test datasets from cancer driver datasets using RFE.

| Classifier | Sensitivity | Specificity | Accuracy | AUC | MCC |
|---|---|---|---|---|---|
| AB | 92.44 | 91.94 | 92.2 | 0.97 | 0.84 |
| DT | 83.11 | 69.67 | 76.61 | 0.85 | 0.53 |
| ET | 95.56 | 97.63 | 96.56 | 0.99 | 0.93 |
| GNB | 79.11 | 74.41 | 76.83 | 0.86 | 0.54 |
| KNN | 79.56 | 96.21 | 87.61 | 0.97 | 0.77 |
| LR | 81.78 | 72.99 | 77.52 | 0.85 | 0.55 |
| MLP | 91.11 | 95.26 | 93.12 | 0.97 | 0.86 |
| RF | 94.67 | 95.73 | 95.18 | 0.99 | 0.9 |
| SVC | 94.67 | 96.68 | 95.64 | 0.98 | 0.9 |
| XGB | 94.67 | 96.21 | 95.41 | 0.99 | 0.91 |

**Table S6**. The performance of gastric cancer classification was developed using 50 features where sensitivity, specificity, and accuracy were measured in percentage using test datasets from cancer driver datasets using MI.

| Classifier | Sensitivity | Specificity | Accuracy | AUC | MCC |
|---|---|---|---|---|---|
| AB | 92 | 93.36 | 92.66 | 0.97 | 0.85 |
| DT | 88 | 69.19 | 78.9 | 0.86 | 0.58 |
| ET | 96.44 | 94.79 | 95.64 | 0.99 | 0.91 |
| GNB | 73.78 | 73.46 | 73.62 | 0.84 | 0.47 |
| KNN | 72.89 | 95.26 | 83.72 | 0.96 | 0.7 |
| LR | 78.67 | 71.09 | 75 | 0.84 | 0.5 |
| MLP | 89.78 | 94.31 | 91.97 | 0.96 | 0.84 |
| RF | 92.89 | 93.84 | 93.35 | 0.98 | 0.87 |
| SVC | 96 | 95.73 | 95.87 | 0.98 | 0.91 |
| XGB | 93.33 | 94.79 | 94.04 | 0.98 | 0.88 |

**Table S7**. The performance of gastric cancer classification was developed using 50 features where sensitivity, specificity, and accuracy were measured in percentage using test datasets from cancer driver datasets using SelectFromModel.

| Classifier | Sensitivity | Specificity | Accuracy | AUC | MCC |
|---|---|---|---|---|---|
| AB | 92 | 94.31 | 93.12 | 0.97 | 0.86 |
| DT | 88.44 | 67.3 | 78.21 | 0.83 | 0.57 |
| ET | 95.11 | 96.68 | 95.87 | 0.99 | 0.92 |
| GNB | 77.78 | 78.67 | 78.21 | 0.86 | 0.56 |
| KNN | 76 | 95.73 | 85.55 | 0.96 | 0.73 |
| LR | 80.44 | 72.04 | 76.38 | 0.86 | 0.53 |
| MLP | 92.89 | 97.16 | 94.95 | 0.97 | 0.9 |
| RF | 95.11 | 94.31 | 94.72 | 0.99 | 0.89 |
| SVC | 92 | 94.79 | 93.35 | 0.98 | 0.91 |
| XGB | 90.22 | 96.21 | 93.12 | 0.99 | 0.86 |

**Table S8**. The performance of gastric cancer classification was developed using 13 features where sensitivity, specificity, and accuracy were measured in percentage using test datasets from the complete dataset.

| Classifier | Sensitivity | Specificity | Accuracy | AUC | MCC |
|---|---|---|---|---|---|
| AB | 88.44 | 93.36 | 90.83 | 0.97 | 0.82 |
| DT | 82.67 | 81.52 | 82.11 | 0.88 | 0.64 |
| ET | 95.11 | 93.36 | 94.27 | 0.99 | 0.89 |
| GNB | 83.56 | 70.62 | 77.29 | 0.82 | 0.55 |
| KNN | 80 | 91 | 85.32 | 0.95 | 0.71 |
| LR | 77.33 | 72.04 | 74.77 | 0.81 | 0.49 |
| MLP | 90.67 | 93.36 | 91.97 | 0.97 | 0.84 |
| RF | 95.11 | 95.26 | 95.18 | 0.99 | 0.9 |
| SVC | 92 | 89.1 | 90.6 | 0.97 | 0.83 |
| XGB | 91.56 | 95.73 | 93.58 | 0.98 | 0.87 |

**Table S9**. The performance of gastric cancer classification was developed using 17 features where sensitivity, specificity, and accuracy were measured in percentage using test datasets from the cancer driver dataset.

| Classifier | Sensitivity | Specificity | Accuracy | AUC | MCC |
|---|---|---|---|---|---|
| AB | 91.11 | 91.94 | 91.51 | 0.96 | 0.83 |
| DT | 83.11 | 66.82 | 75.23 | 0.79 | 0.51 |
| ET | 94.67 | 92.89 | 93.81 | 0.98 | 0.88 |
| GNB | 76 | 78.2 | 77.06 | 0.84 | 0.54 |
| KNN | 81.78 | 94.31 | 87.84 | 0.96 | 0.76 |
| LR | 78.67 | 73.93 | 76.38 | 0.84 | 0.53 |
| MLP | 88.44 | 89.57 | 88.99 | 0.96 | 0.78 |
| RF | 94.67 | 95.26 | 94.95 | 0.98 | 0.9 |
| SVC | 92.89 | 91.94 | 92.43 | 0.96 | 0.8 |
| XGB | 93.33 | 93.36 | 93.35 | 0.98 | 0.87 |

**Table S10.** 14 prognostic genes with their gene symbol, logFC, P.value, adj.P.value, regulation, and AUC.

| Gene | logFC | P.Value | adj.P.Val | Regulation | AUC |
|---|---|---|---|---|---|
| PLCXD3 | -2.68512 | 1.27E-105 | 8.54E-102 | DOWN | 0.770213 |
| LIPG | 2.291105 | 4.17E-53 | 8.33E-51 | UP | 0.736612 |
| BGN | 2.134639 | 2.64E-52 | 4.93E-50 | UP | 0.729981 |
| PCOLCE2 | -1.90391 | 1.03E-34 | 4.08E-33 | DOWN | 0.66739 |
| SPARC | 1.961244 | 1.88E-23 | 2.44E-22 | UP | 0.707634 |
| AGT | 1.623641 | 6.56E-23 | 8.10E-22 | UP | 0.65836 |
| BEX2 | -1.24699 | 1.81E-18 | 1.47E-17 | DOWN | 0.6647 |
| COL11A1 | 3.123094 | 5.32E-81 | 6.31E-78 | UP | 0.663581 |
| IRS4 | -1.70751 | 1.42E-47 | 1.76E-45 | DOWN | 0.672029 |
| CDH11 | 1.554493 | 7.12E-28 | 1.44E-26 | UP | 0.645533 |
| MYB | 1.692607 | 6.50E-27 | 1.18E-25 | UP | 0.67689 |
| PDGFRB | 1.414139 | 1.52E-25 | 2.41E-24 | UP | 0.668767 |
| COL5A1 | 1.395381 | 7.12E-22 | 7.93E-21 | UP | 0.631903 |
| COL3A1 | 1.556473 | 5.62E-13 | 2.59E-12 | UP | 0.647597 |

***Supplementary Figures***

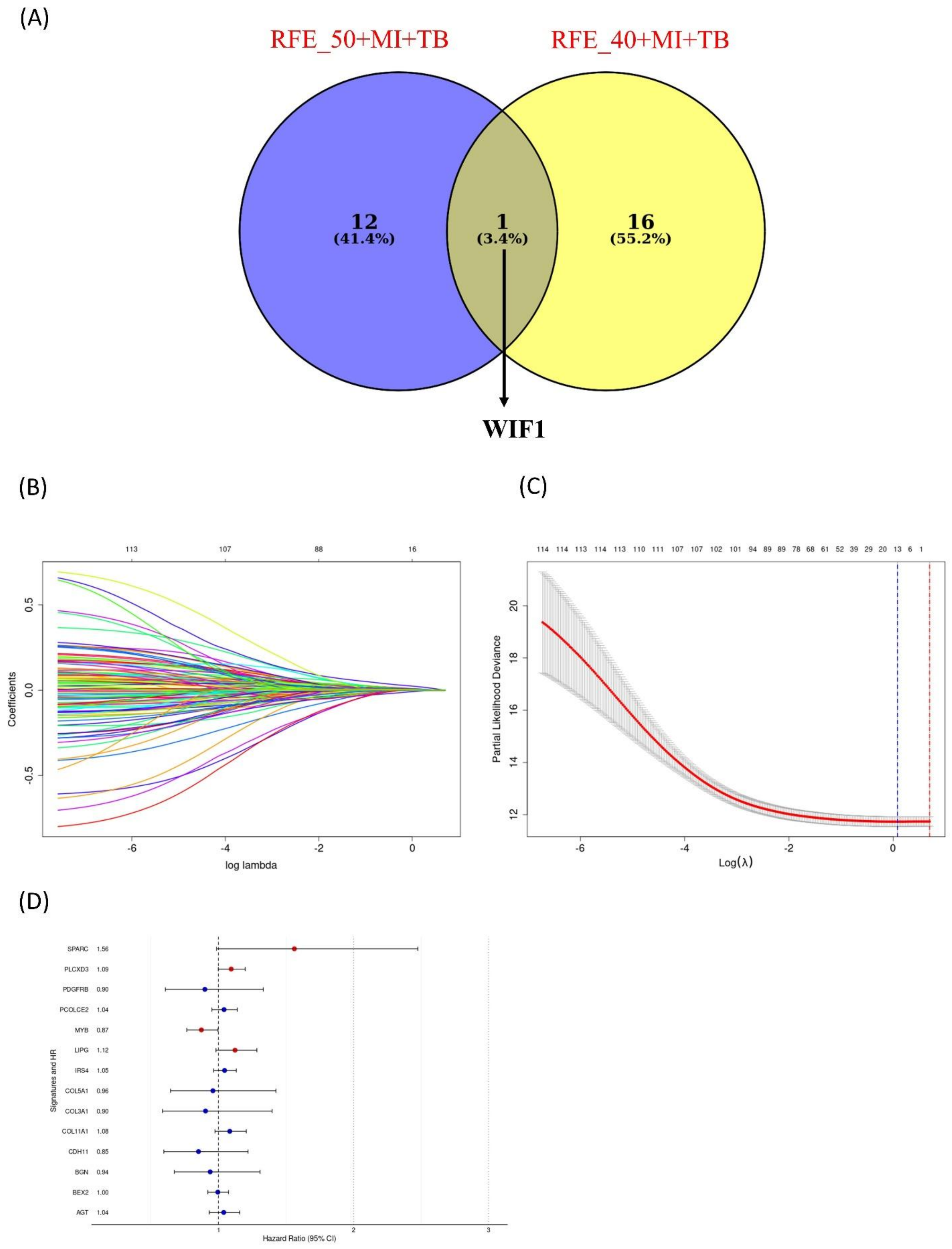


**Figure S1:** (A) A Venn diagram showing WIF1 as only the shared feature between 13 genes from the complete dataset and 17 genes from the cancer driver genes. (B) Key gene selection using Lasso penalized Cox regression. (C) Coefficient distribution map for a logarithmic sequence of λ. (D) Forest plot of the univariate Cox model, displaying the hazard ratio (HR) as a relative prognostic indicator for gastric cancer patients. The red color highlights statistical significance, with p-values < 0.05.

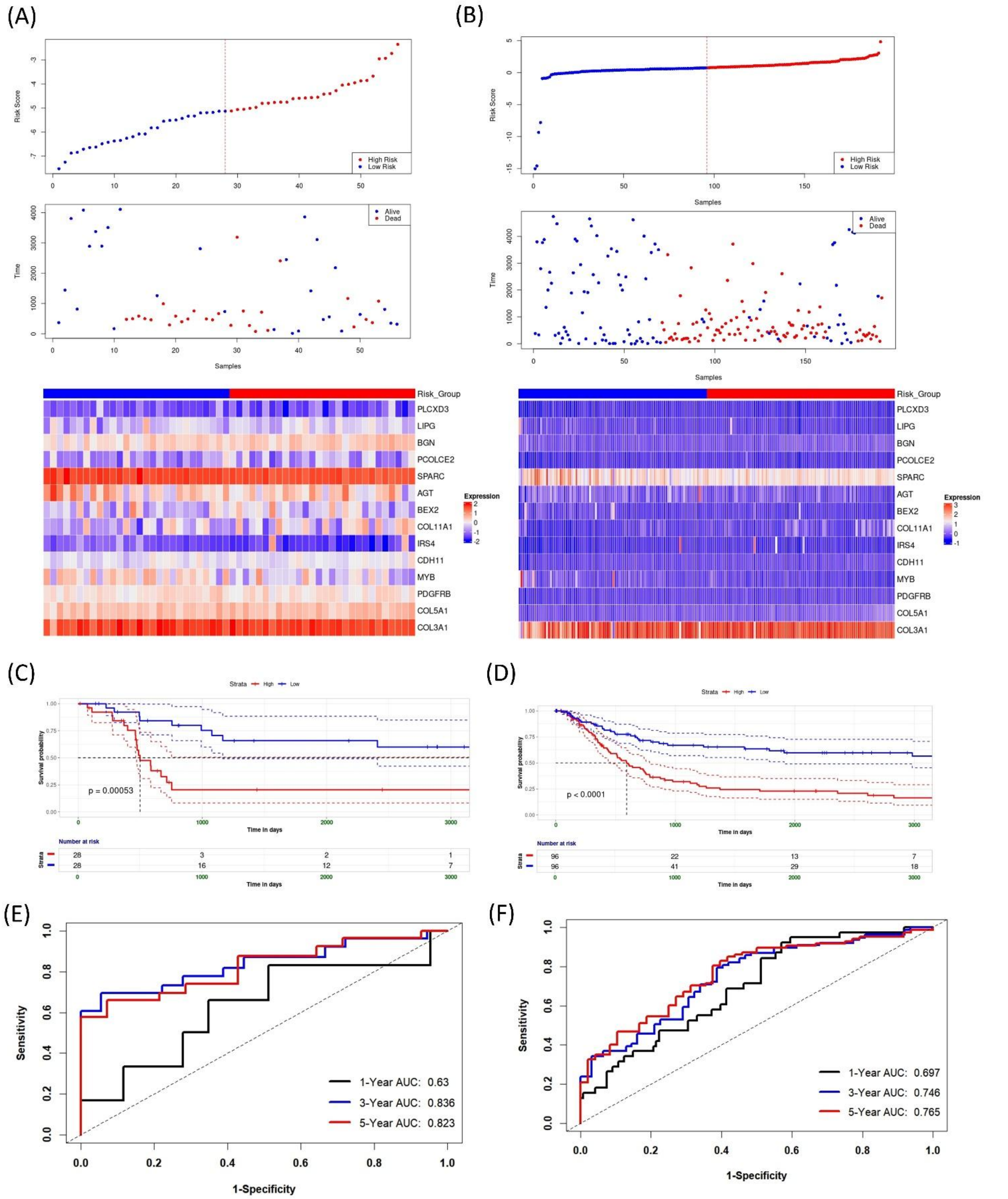


**Figure S2:** The performance of the fourteen-gene model in GSE34942 and GSE15459 is shown in panels (A-B) These panels display the distribution of risk scores between high and low-risk groups, the density distribution of risk scores in deceased and surviving patients, and heat maps illustrating the expression levels of the fourteen genes. (C-D) Survival analysis (E-F) ROC curve of a risk model for predicting survival for both cohorts.
